# Increased power from bacterial genome-wide association conditional on known effects identifies *Neisseria gonorrhoeae* macrolide resistance mutations in the 50S ribosomal protein L4

**DOI:** 10.1101/2020.03.24.006650

**Authors:** Kevin C Ma, Tatum D Mortimer, Marissa A Duckett, Allison L Hicks, Nicole E Wheeler, Leonor Sánchez-Busó, Yonatan H Grad

**Affiliations:** Department of Immunology and Infectious Diseases, Harvard T.H. Chan School of Public Health, Boston, USA; Centre for Genomic Pathogen Surveillance, Wellcome Sanger Institute, Wellcome Genome Campus, Hinxton, Cambridgeshire, United Kingdom; Division of Infectious Diseases, Brigham and Women’s Hospital and Harvard Medical School, Boston, USA

## Abstract

The emergence of resistance to azithromycin complicates treatment of *N. gonorrhoeae*, the etiologic agent of gonorrhea. Population genomic analyses of clinical isolates have demonstrated that some azithromycin resistance remains unexplained after accounting for the contributions of known resistance mutations in the 23S rRNA and the MtrCDE efflux pump. Bacterial genome-wide association studies (GWAS) offer a promising approach for identifying novel resistance genes but must adequately address the challenge of controlling for genetic confounders while maintaining power to detect variants with lower effect sizes. Compared to a standard univariate GWAS, conducting GWAS conditioned on known resistance mutations with high effect sizes substantially reduced the number of variants that reached genome-wide significance and identified a G70D mutation in the 50S ribosomal protein L4 (encoded by the gene *rplD*) as significantly associated with increased azithromycin minimum inhibitory concentrations (β = 1.03, 95% CI [0.76, 1.30]). The role and prevalence of these *rplD* mutations in conferring macrolide resistance in *N. gonorrhoeae* had been unclear. Here, we experimentally confirmed our GWAS results, identified other resistance-associated mutations in RplD, and showed that in total these RplD binding site mutations are prevalent (present in 5.42% of 4850 isolates) and geographically and temporally widespread (identified in 21/65 countries across two decades). Overall, our findings demonstrate the utility of conditional associations for improving the performance of microbial GWAS and advance our understanding of the genetic basis of macrolide resistance in a prevalent multidrug-resistant pathogen.

## Introduction

Increasing antibiotic resistance in *Neisseria gonorrhoeae*, the causative agent of the sexually transmitted disease gonorrhea, threatens effective control of this prevalent pathogen [1-3]. Current empiric antibiotic therapy in the US comprises a combination of the cephalosporin ceftriaxone and the macrolide azithromycin, but increasing prevalence of azithromycin resistance has led some countries, such as the UK, to instead recommend ceftriaxone monotherapy [4]. Rapid genotypic diagnostics for antimicrobial susceptibility have been proposed as a platform to tailor therapy and to extend the clinically useful lifespan of anti-gonococcal antibiotics [5, 6]. These rapid diagnostics rest on robust genotype-to-phenotype predictions. For some antibiotics, such as ciprofloxacin, resistance is predictable by target site mutations in a single gene, *gyrA* [3, 5]. However, recent efforts to predict azithromycin minimum inhibitory concentrations (MICs) using regression-based or machine-learning approaches have indicated that a substantial fraction of phenotypic resistance is unexplained, particularly among strains with lower-level resistance [3, 7, 8]. An improved understanding of the genetic mechanisms and evolutionary pathways to macrolide resistance will therefore be critical for informing the development of diagnostics.

Macrolides function by binding to the 50S ribosome and inhibiting protein synthesis [9]. Increased resistance can occur in *N. gonorrhoeae* through target site modification, primarily via 23S rRNA mutations C2611T [10] and A2059G [11], and though efflux pump upregulation. The main efflux pump associated with antibiotic resistance in the gonococcus is the Mtr efflux pump, comprising a tripartite complex encoded by the *mtrCDE* operon under the regulation of the MtrR repressor and the MtrA activator [1, 12-17]. Active site or frameshift mutations in the coding sequence of *mtrR* and promoter mutations in the *mtrR* promoter upregulate *mtrCDE* and result in increased macrolide resistance [1, 18]. Mosaic sequences originating from recombination with homologs from commensal *Neisseria* donors can also result in structural changes to *mtrD* and increased expression of *mtrCDE*, which synergistically act to confer resistance [19, 20].

Here, we used genome-wide association on a global meta-analysis dataset to identify additional genetic variants that confer increased azithromycin resistance in *N. gonorrhoeae.* We found that conventional single-locus bacterial GWAS approaches that univariately test genetic variants resulted in confounded results and reduced power. Conducting GWAS conditional on known resistance mutations in 23S rRNA reduced linkage-mediated confounding and increased power to recover known and candidate mutations associated with lower-level resistance. We experimentally validated one such mutation in the 50S ribosomal protein RplD and identified other rare RplD variants associated with resistance, highlighting the ability of conditional bacterial GWAS to identify causal genes for polygenic microbial phenotypes.

## Results

We previously conducted a linear mixed model GWAS on continuous azithromycin MICs in 4535 *N. gonorrhoeae* isolates where we observed highly significant unitigs (i.e., genetic variants generated from *de novo* assemblies) mapping to the 23S rRNA, associated with increased resistance, and to the efflux pump gene *mtrC*, associated with increased susceptibility and cervical infections [7]. We re-analyzed the GWAS results focusing on the remaining significant variants, which were closer in significance to the Bonferonni-corrected *p*-value threshold of 2.97×10^−7^. Numerous variants were significantly associated with increased MICs, many of which mapped to genes (e.g., *hprA, ydfG*, and *efeB*) that had not previously been implicated in macrolide resistance in *Neisseria* (Supplementary Table 1). While these signals could represent novel causal resistance genes, we hypothesized that at least some of these variants had been spuriously driven to association via genetic linkage with the highly penetrant (A2059G: β, or effect size, = 7.21, 95% CI [6.52, 7.90]; C2611T: β = 3.62, 95% CI [3.42, 3.82]) and population-stratified 23S rRNA resistance mutations (Supplementary Figure 1). Supporting this hypothesis, *r*^2^ – a measure of linkage ranging from 0 to 1 – between significant variants and 23S rRNA resistance mutations exhibited a bimodal distribution with a peak at 0.84 and at 0.04 (Supplementary Figure 2). The three most significant variants after the 23S rRNA substitutions and *mtrC* deletion mapping to *hprA*, WHO_F.1254, and *ydfG* had elevated *r*^2^ values of 0.16, 0.82, and 0.80 respectively; all three variants demonstrated clear phylogenetic overlap with 23S rRNA mutations (Supplementary Figure 1). We additionally did not observe unitigs associated with experimentally validated resistance mutations in the *mtrR* promoter [14] or the *mtrCDE* mosaic alleles [19, 20], suggesting decreased power to detect known causal variants with lower effect sizes.

To control for the confounding effect of the 23S rRNA mutations, we conducted a conditional GWAS by incorporating additional covariates in our linear mixed model encoding the number of copies of the resistance-conferring 23S rRNA substitutions C2611T and A2059G. We also conditioned on isolate dataset of origin to address potential spurious hits arising from study-specific sequencing methodologies. After conditioning, the previously significant genes linked to 23S rRNA (*r*^2^ > 0.80) decreased below the significance threshold, indicating that they were indeed driven to significance by genetic linkage (Figure 1, Supplementary Table 2). The most significant variants after the previously reported *mtrC* indel [7] mapped to the *mtrR* promoter (β, or effect size, = –0.79, 95% CI [−0.62, −0.96]; *p*-value = 1.62×10^−18^), encoding the *mtrR* promoter 1 bp deletion [21], and to *mtrC* (β = 1.21, 95% CI [0.92, 1.50]; *p*-value = 9.17×10^−16^), in linkage with mosaic *mtr* alleles [19, 20]. The increased significance of these known efflux pump resistance mutations suggested improved power to recover causal genes with lower effects. Conditioning on dataset did not substantially affect these results but helped to remove other spurious variants arising due to study-specific biases (Supplementary Figure 3, Supplementary Table 3).

**Figure 1.**
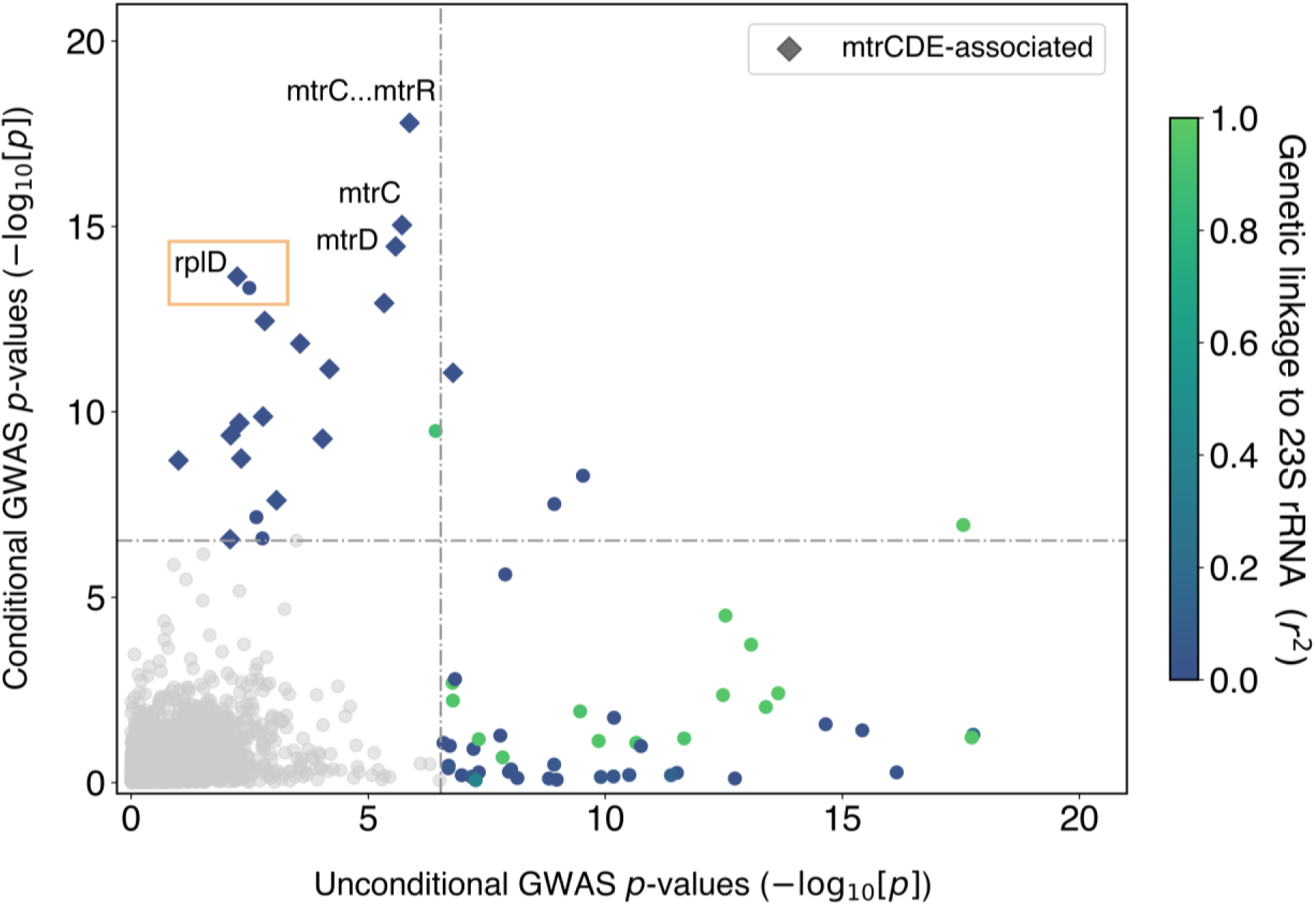
GWAS conditional on 23S rRNA mutations and dataset demonstrates decreased confounding due to genetic linkage and increased power to recover known and candidate lower-level resistance alleles. Genetic linkage measured by *r*^*2*^ to 23S rRNA mutations A2059G and C2611T is colored as indicated on the right. Variants associated with previously experimentally verified resistance mechanisms in the *mtrR* and *mtrCDE* promoters and coding regions are denoted in the legend. Bonferroni thresholds for both GWASes are depicted using a dashed line at 2.97×10^−7^. Plot axes are limited to highlight variants associated with lower-level resistance; as a result, the highly significant 23S rRNA substitutions and *mtrC* indel mutations are not shown.

A glycine to glutamic acid substitution at site 70 of the 50S ribosomal protein L4 (RplD) was significantly associated with increased azithromycin MICs after conducting the conditional GWAS (β = 1.03, 95% CI [0.76, 1.30]; *p*-value = 4.56×10^−14^) (Figure 1, Supplementary Table 2). Structural analysis of the *Thermus thermophilus* 50S ribosome complexed with azithromycin suggests that this amino acid is an important residue in macrolide binding (Supplemental Figure 4), and RplD substitutions at this binding site modulate macrolide resistance in other bacteria [22, 23]. This substitution has previously been observed rarely in gonococcus and the association with binarized azithromycin resistance was non-significant [3, 24, 25]; as a result, the role of RplD mutations in conferring macrolide resistance was unclear. To assess the contribution of RplD mutations to continuous azithromycin MIC levels, we modeled MICs using a linear regression framework with known genetic resistance determinants as predictors [7, 26]. Compared to this baseline model, inclusion of the RplD G70D mutation decreased the number of strains with unexplained MIC variation (defined as absolute model error greater than one MIC dilution) from 1430 to 1333, improved adjusted *R*^2^ from 0.721 to 0.734, and significantly improved model fit (*p*-value < 2.2×10^−16^; Likelihood-ratio X^2^ test for nested models). These results indicate that RplD G70D is a strong candidate for addressing a portion of the unexplained azithromycin resistance in *N. gonorrhoeae*.

We next assessed population-wide prevalence and diversity of RplD-azithromycin binding site mutations. The RplD G70D mutation was present in 231 out of 4850 isolates (4.76%) with multiple introductions observed across varied genetic backgrounds (Figure 2). An additional 34 isolates contained mutations at amino acids 68 (G68D, G68C), 69 (T69I), and 70 (G70S, G70A, G70R, G70duplication) (Figure 3). These other putative RplD binding site mutations were associated with significantly higher azithromycin MICs compared to both RplD G70D and RplD wild-type strains, indicating multiple avenues for disruption of macrolide binding (Figure 3). Strains with RplD binding site mutations were identified from 21 countries from 1993 to 2015 with prevalence reaching over 10% in some datasets (New York City 2011-2015 [Mortimer *et al.*, 2020] and Japan 1996-2015 [27]; Supplementary Table 4 and [28]), in line with sustained transmission of RplD G70D strains (Figure 2). Our results suggest that macrolide binding to the 50S ribosome can be disrupted via multiple mutations and that these mutations are widespread contributors to azithromycin resistance in some populations.

**Figure 2.**
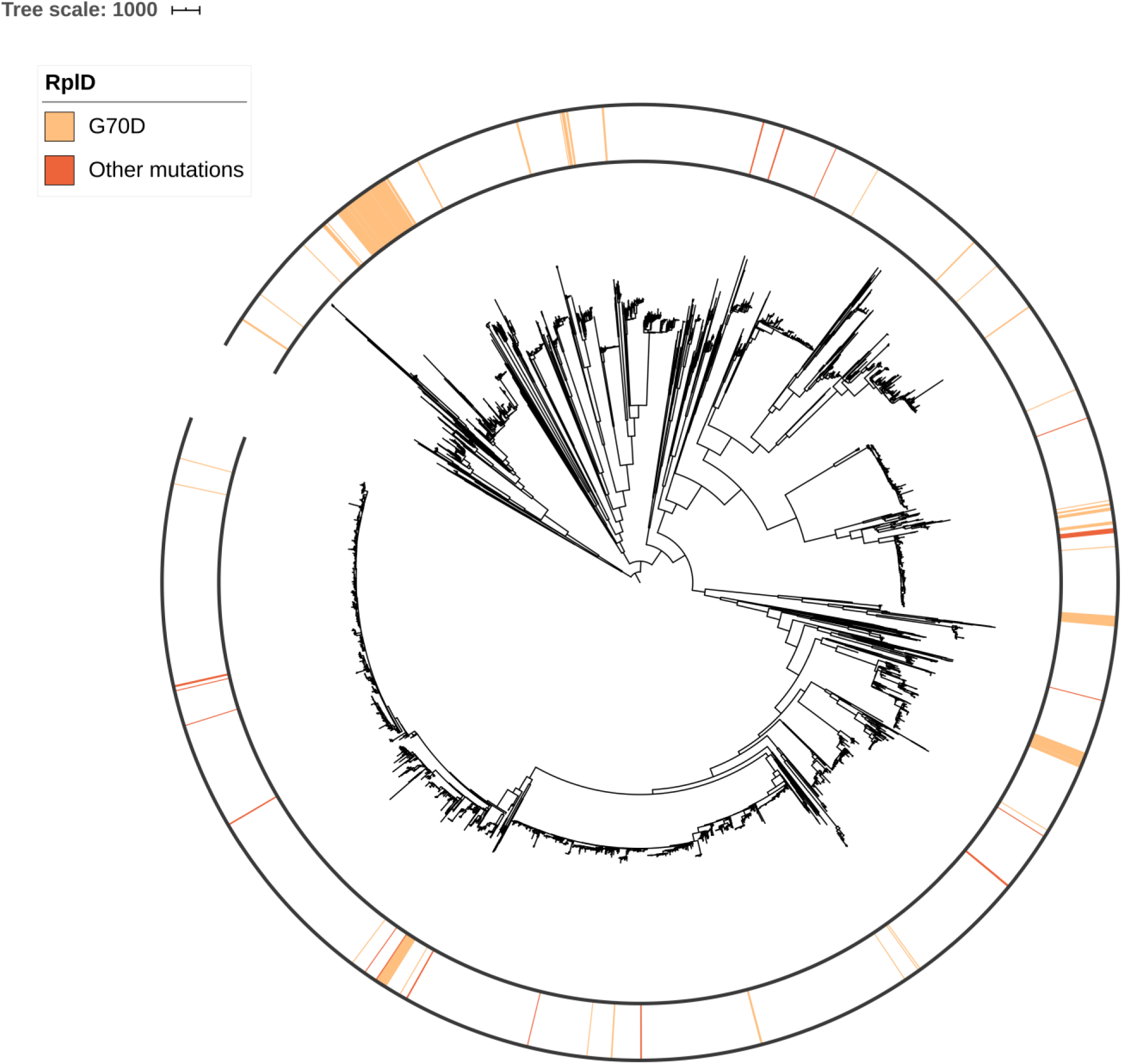
Population structure of RplD binding site mutations in a global meta-analysis dataset of *N. gonorrhoeae*. A midpoint rooted recombination-corrected maximum likelihood phylogeny of 4882 genomes based on 68697 SNPs non-recombinant from [7] was annotated with the presence of RplD binding site mutations Branch length represents total number of substitutions after removal of predicted recombination.

**Figure 3.**
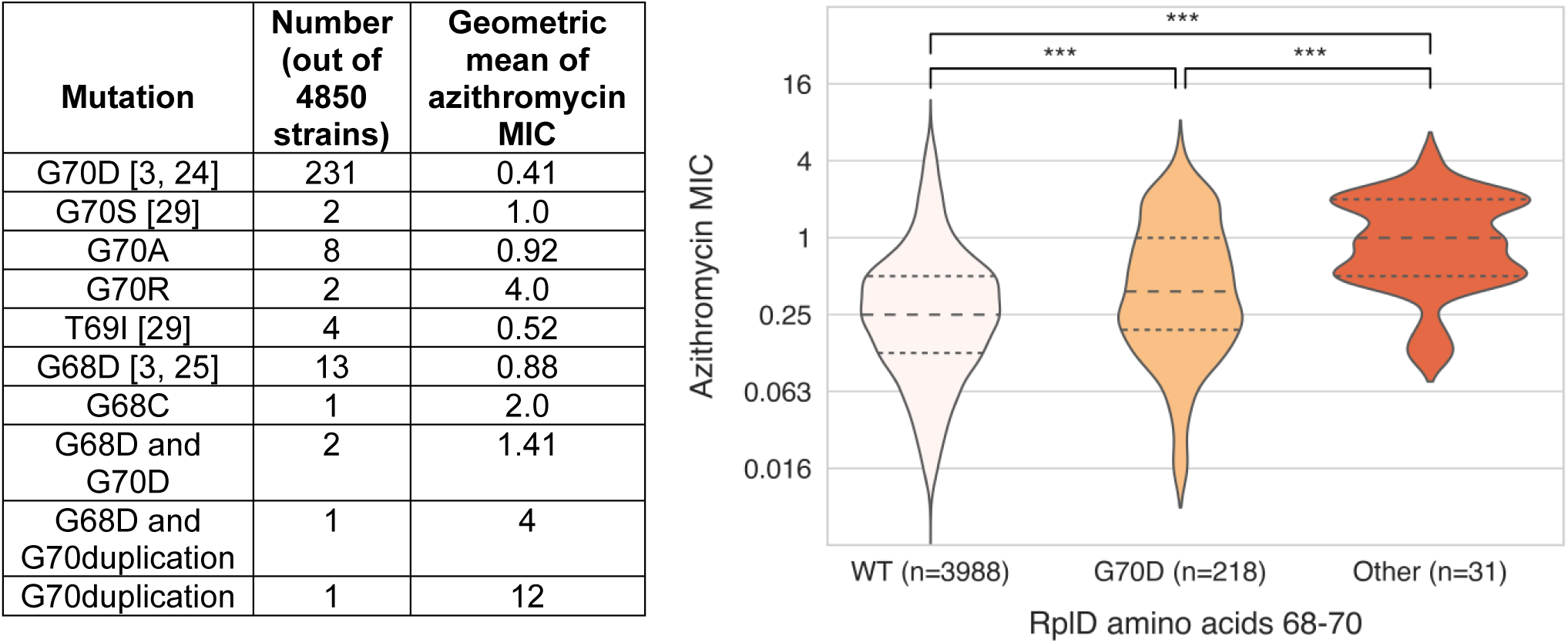
RplD amino acid diversity at positions 68 through 70 and corresponding geometric mean (left) and distribution (right) of azithromycin MICs. Previously reported mutations are cited with the first reporting publications. Violin plots and statistical analyses were limited to isolates with MICs < 8 to exclude isolates with 23S rRNA mutations. Quartiles within violin plots are depicted using dotted lines. Statistical significance between RplD variants and RplD wildtype MIC distributions was assessed by Mann-Whitney U Test: * p<0.05, ** p<0.01, and *** p<0.001.

To experimentally verify that RplD G70D contributes to macrolide resistance, we constructed two isogenic strains (C5 and E9) with the G70D substitution and tested for MIC differences across a panel of macrolides. Azithromycin and erythromycin MICs increased by three-fold, and clarithromycin MICs increased by six-fold on average in the G70D strains compared to isogenic wild-type strains (Table 1). The estimate from our linear model for the azithromycin MIC of a strain that contains the RplD G70D mutation and no other resistance mutations was 0.252, which is in line with the experimental results. Macrolide resistance has been associated with a fitness cost in other species [30], prompting us to measure the *in vitro* growth dynamics of the RplD G70D strain. Time-course growth curves of the wild-type strain and isogenic G70D strain E9 were similar (Supplementary Figure 5) with overlapping estimates of doubling times: 28Bl doubling time = 1.756 hours, 95% CI [1.663, 1.861] versus 28Bl RplD^G70D^ (E9) doubling time = 1.787 hours, 95% CI [1.671, 1.920] (Supplementary Table 5). These results confirm the role of RplD G70D in mediating macrolide resistance and indicate a lack of severe associated *in vitro* fitness costs.

**Table 1.**
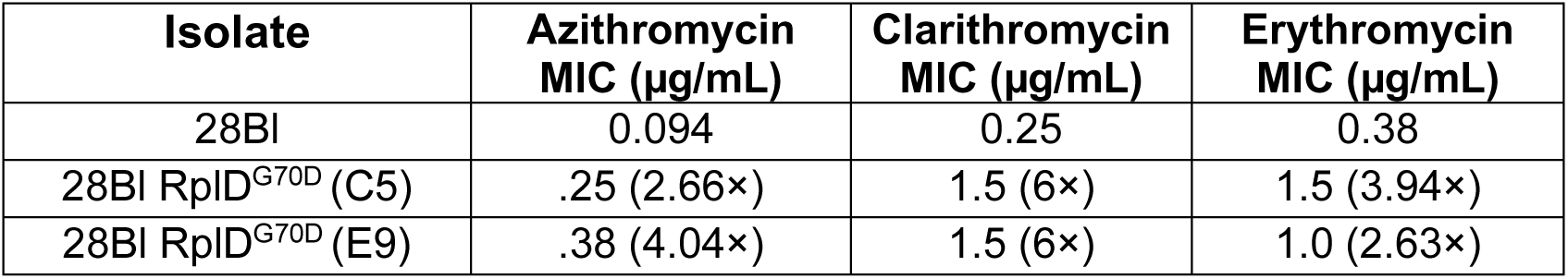
Macrolide MICs of laboratory strain 28Bl and two isogenic derivatives confirms increased macrolide resistance conferred by RplD G70D. Fold change relative to baseline is shown in parentheses. MICs were measured using Etest strips placed onto GCB agar plates supplemented with 1% IsoVitaleX.

## Discussion

Azithromycin resistance in *N. gonorrhoeae* is a polygenic trait involving contributions from mutations in different 50S ribosomal components, up- and down-regulation of efflux pump activity, and additional unknown factors. Genome-wide association methods offer one approach for uncovering the genotypic basis of unexplained resistance in clinical isolates, but novel causal genes associated with lower effects have been difficult to identify with traditional microbial GWAS approaches [23]. Our results indicate that extending the GWAS linear mixed model to incorporate known causal genetic variants could address some of these challenges, particularly when known genes exhibit strong penetrance and population stratification, obfuscating signals with lower effects. After conducting conditional GWAS on azithromycin MICs, we observed a reduction in spurious results attributable to genetic linkage with known high level resistance mutations in the 23S rRNA and an increase in power to recover secondary resistance mutations in the MtrCDE efflux pump. We also identified a resistance-associated mutation in the macrolide binding site of 50S ribosomal protein RplD as significant only after conditioning. While the improvements in GWAS performance suggested by these empirical results will need to be further validated on other bacterial species and through simulations [31], they are in line with studies of multi-locus methods in the human GWAS field [32, 33] and complementary methods using whole-genome elastic nets for microbial genome data [31, 34].

The role of RplD G70D mutations in conferring azithromycin resistance has previously been unclear, in part because of its lower effect size relative to 23S rRNA mutations. The G70D mutation was first observed in isolates from France 2013-2014 [24] and in the US Centers for Disease Control Gonococcal Isolate Surveillance Program (CDC GISP) surveillance isolates from 2000-2013 [3], and a related G68D mutation was described in the GISP collection and in European isolates from 2009-2014 [25]. However, these analyses reported no clear association with binarized resistance levels. Follow up studies in the US, Eastern China, and a historical Danish collection also reported strains with the G70D mutation [29, 35, 36], but other surveillance datasets from Canada, Switzerland, and Nanjing did not [10, 37-39], indicating geography-specific circulation. As a result of this ambiguity, previous studies modeling phenotypic azithromycin resistance from genotype did not include RplD mutations [26, 40].

Here, we provided confirmatory evidence that the RplD G70D mutation increases macrolide MICs several-fold, in line with the GWAS analyses. While RplD G70D mutations on their own are not predicted to confer resistance levels above the clinical CLSI non-susceptibility threshold of 1.0 µg/mL, there is growing appreciation of the role that sub-breakpoint increases in resistance can play in mediating treatment failure [41]. For example, treatment failures in Japan after a 2 g azithromycin dose were associated with MICs as low as 0.5 µg/mL [42], and treatment failures in several case studies of patients treated with a 1 g azithromycin dose were associated with MICs of 0.125 to 0.25 µg/mL [43]. Low level azithromycin resistance may also serve as a stepping stone to higher level resistance, as suggested by an analysis of an outbreak of a high level azithromycin resistant *N. gonorrhoeae* lineage in the UK [44].

We also observed multiple previously undescribed in the RplD macrolide binding site associated with even higher MICs than the G70D mutation. The transmission of these isolates has been relatively limited, potentially due to increased fitness costs commensurate with increased resistance. In contrast, several lines of evidence suggest that the G70D mutation carries a relatively minimal fitness cost. Time-course growth experiments indicated that the RplD G70D isogenic pair of strains have similar doubling times, and phylogenetic analyses suggest multiple acquisitions of G70D in distinct genetic backgrounds with a lineage in NYC showing evidence of sustained transmission. As macrolide use continues to select for increased resistance in *N. gonorrhoeae*, both the RplD G70D and rarer binding site mutations should be targets for surveillance in future whole-genome sequencing studies.

In summary, by reducing genetic confounders and amplifying true signals through bacterial GWAS conditional on known effects, we identified and experimentally characterized mutations in the 50S ribosome that contribute to increased macrolide resistance in *N. gonorrhoeae*.

## Methods

### Genomics and GWAS

We conducted whole-genome sequencing assembly, resistance allele calling, phylogenetic inference, genome-wide association, and significant unitig mapping using methods from a prior GWAS [7]. Briefly, we created a recombination-corrected phylogeny by running Gubbins (version 2.3.4) [45] on an alignment of pseudogenomes generated from filtered SNPs from Pilon (version 1.16) [46] after mapping reads in BWA-MEM (version 0.7.17-r1188) [47] to a reference genome (RefSeq accession: NC_011035.1). To conduct the GWAS in Pyseer (version 1.2.0) [48], unitigs were generated from GATB using SPAdes (version 3.12.0) [49] *de novo* assembled genomes, and a population structure matrix was generated from the Gubbins phylogeny for the linear mixed model. Isolates included in this study are listed in Supplementary Table 6. As in the prior study, azithromycin MICs prior to 2005 from the CDC GISP dataset were doubled to account for an MIC protocol testing change [50]. We conducted conditional GWAS in Pyseer (version 1.2.0) [48] by including additional columns in the covariate file encoding 23S rRNA mutations and including flags --covariates and --use-covariates. All phylogenies and annotation rings were visualized in iTOL (version 5.5) [51].

We assessed genetic linkage by calculating *r*^2^, or the squared correlation coefficient between two variants defined as *r*^2^ = (*p*_*ij*_ – *p*_*i*_*p*_*j*_)^2^ / (*p*_*i*_ (1 – *p*_*i*_) *p*_*j*_ (1 – *p*_*j*_)), where *p*_*i*_ is the proportion of strains with variant *i, p*_*j*_ is the proportion of strains with variant *j*, and *p*_*ij*_ is the proportion of strains with both variants [52, 53]. For a given GWAS variant, we calculated *r*^2^ between that variant and the significant unitig from the GWAS mapping to 23S rRNA C2611T. We repeated the calculation for the same variant but with the unitig mapping to 23S rRNA A2059G, and took the maximum *r*^2^ value from the two calculations.

Azithromycin log-transformed MICs were modeled using a panel of resistance markers [7] including pairwise interactions and country of origin in R (version 3.5.1), with and without inclusion of RplD G70D and proximal mutations:

Model 1: Country + (MtrR 39 + MtrR 45 + MtrR LOF + MtrC LOF + MtrR promoter + MtrCDE BAPS + 23S rRNA 2059 + 23S rRNA 2611)^2

Model 2: Country + (MtrR 39 + MtrR 45 + MtrR LOF + MtrC LOF + MtrR promoter + MtrCDE BAPS + 23S rRNA 2059 + 23S rRNA 2611 + RplD G70D + RplD other 68-70 mutations)^2

Model fit was assessed using Anova for likelihood-ratio tests for nested models in R (version 3.5.1). BAPS groups for MtrCDE were called as previously described using FastBAPS (version 1.0.0) [7].

### Diversity of RplD macrolide binding site mutations

We ran BLASTn (version 2.6.0) [54] on the *de novo* assemblies using a query *rplD* sequence from FA1090 (Genbank accession: NC_002946.2). *rplD* sequences were aligned using MAFFT (version 7.450) [55]. Binding site mutations were identified after *in silico* translation of nucleotide alignments in Geneious Prime (version 2019.2.1, https://www.geneious.com). Subsequent analyses identifying prevalence, geometric mean azithromycin MIC, and MIC distribution differences were conducted in Python (version 3.6.5) and R (version 3.5.1).

### Experimental validation

We cultured *N. gonorrhoeae* on GCB agar (Difco) plates supplemented with 1% Kellogg’s supplements (GCBK) at 37°C in a 5% CO2 incubator [56]. We conducted antimicrobial susceptibility testing using Etests (bioMérieux) placed onto GCB agar plates supplemented with 1% IsoVitaleX (Becton Dickinson). We selected laboratory strain 28Bl for construction of isogenic strains and measured its MIC for azithromycin, clarithromycin, and erythromycin [20]. *rplD* encoding the G70D mutation was PCR amplified from RplD G70D isolate GCGS1043 [3] using primers rplD_FWD_DUS (5’ CATGCCGTCTGAACAAGACCCGGGTCGCG 3’) (containing a DUS tag to enhance transformation [57]) and rplD_REV (5’ TTCAGAAACGACAGGCGCC 3’). The resulting ∼1 kb amplicon was spot transformed [56] into 28Bl. We selected for transformants by plating onto GCBK plates with clarithromycin 0.4 μg/mL and erythromycin 0.4 μg/mL. We confirmed via Sanger sequencing that transformants had acquired the RplD G70D mutation and selected one transformant from each selection condition (strain C5 for clarithromycin and strain E9 for erythromycin) for further characterization. We confirmed that for all macrolides used for selection, no spontaneous resistant mutants were observed after conducting control transformations in the absence of GCGS1043 PCR product.

### Growth assays

We streaked 28Bl and 28Bl RplD^G70D^ (E9) onto GCBK plates and grew them overnight for 16 hours at 37°C in a 5% CO2 atmosphere. We prepared 1 L of fresh Graver Wade (GW) media [58] and re-suspended overnight cultures into 1 mL of GW. After normalizing cultures to OD 0.1, we diluted cultures 1:10^5^ and inoculated central wells of a 24-well plate with 1.5 mL GW and cells in triplicate. Edge wells were filled with 1.5 mL water. After growth for 1 hour to acclimate to media conditions, we sampled CFUs every 2 hours for a total of 12 hours. For each timepoint, we aspirated using a P1000 micropipette to dissolve clumps and then plated serial dilutions onto a GCBK plate. We counted CFUs the following day and used GraphPad Prism (version 8.2.0 for Windows, GraphPad Software) to graph the data and estimate exponential phase growth rates following removal of lag phase data points and log-transformation of CFU / mLs.

## Acknowledgements

This work was supported by the NIH/NIAID grant 1R01AI132606-01 and the Smith Family Foundation. TDM is additionally supported by the NIH/NIAID 1 F32 AI145157-01, and KCM is additionally supported by the NSF GRFP. Portions of this research were conducted on the O2 high-performance computing cluster, supported by the Research Computing Group at Harvard Medical School. The authors additionally thank Daniel Rubin, Samantha Palace, Crista Wadsworth, and other members of the Grad Lab for helpful comments during development of the project.

---

**Supplementary Figure 1.**
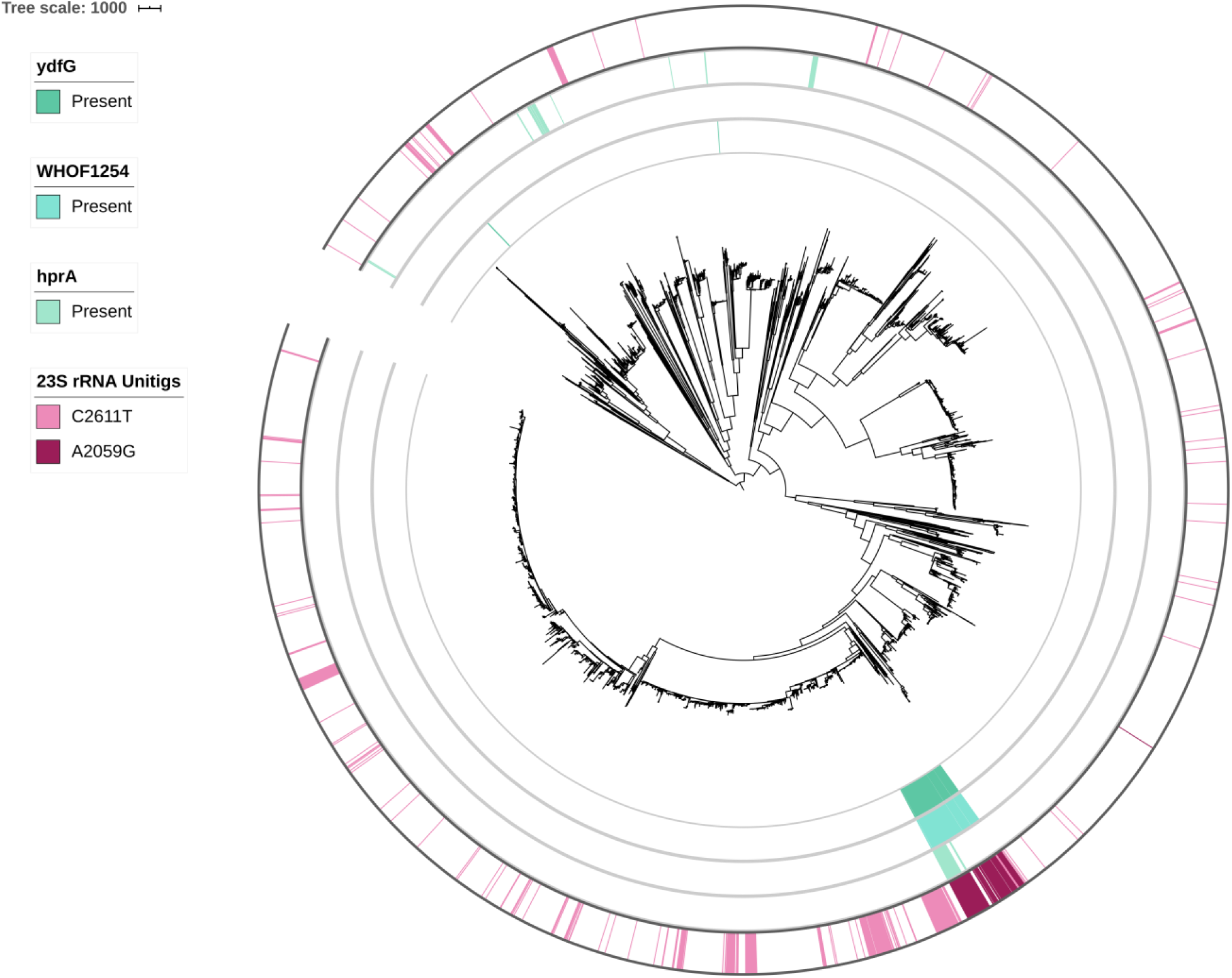
Genetic linkage between significant azithromycin MIC-associated variants in the GWAS. The recombination-corrected phylogeny from Figure 1 was annotated with the presence and absence of significant variants from the GWAS corresponding to 23S rRNA, *hprA*, WHO_F.1254, and *ydfG* (outermost to innermost). Branch length represents total number of substitutions after removal of predicted recombination.

**Supplementary Figure 2.**
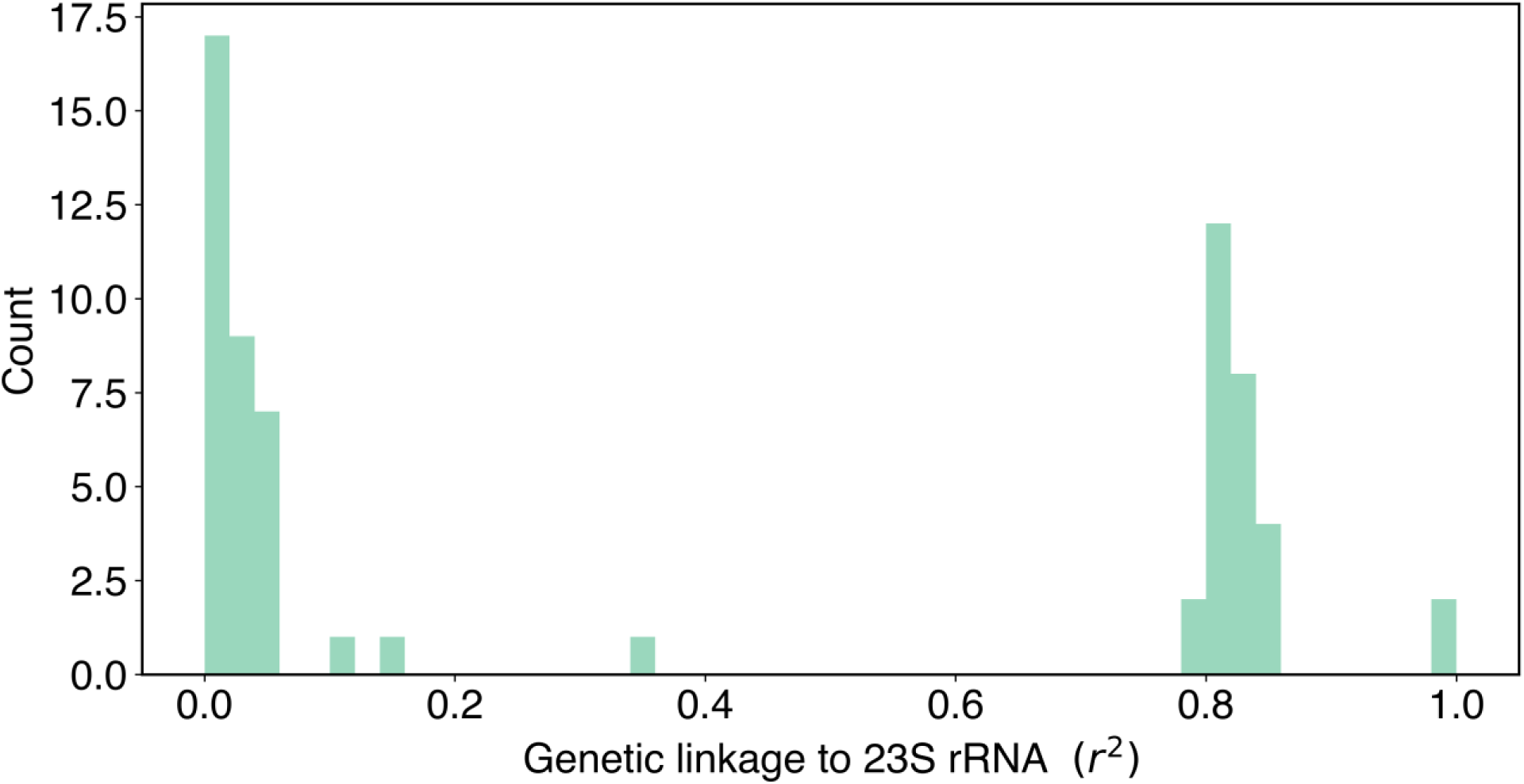
Distribution of *r*^2^ values between significant variants (*p*-value < 2.97×10^−7^) and 23S rRNA-associated unitigs in the single-locus GWAS. Significant variants with high linkage to 23S rRNA are likely to be spurious associations. See methods for details on calculation of *r*^2^.

**Supplementary Figure 3.**
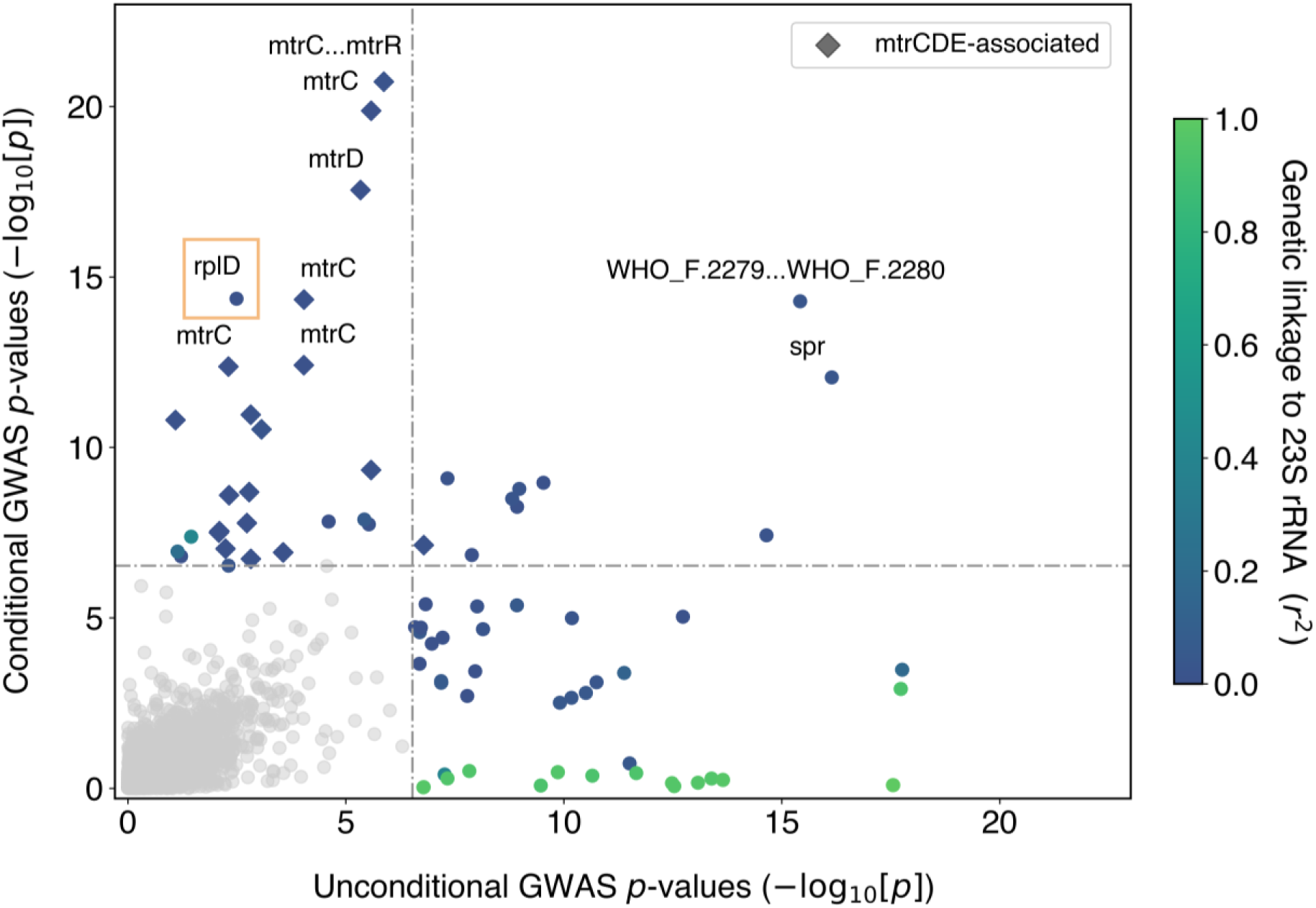
GWAS conditional on 23S rRNA mutations compared to unconditional GWAS results recovers similar results as in Figure 1 but does not control for dataset-specific confounders (*spr* and the intergenic region between genes WHO_F.2279 and 2280). As in Figure 1, genetic linkage measured by *r*^*2*^ to 23S rRNA mutations A2059G and C2611T is colored as indicated on the right. Variants associated with previously experimentally verified resistance mechanisms in the *mtrR* and *mtrCDE* promoters and coding regions are denoted in the legend. Bonferroni thresholds for both GWASes are depicted using a dashed line at 2.97×10^−7^. Plot axes are limited to highlight variants associated with lower-level resistance; as a result, the highly significant 23S rRNA substitutions and *mtrC* indel mutations are not shown.

**Supplementary Figure 4.**
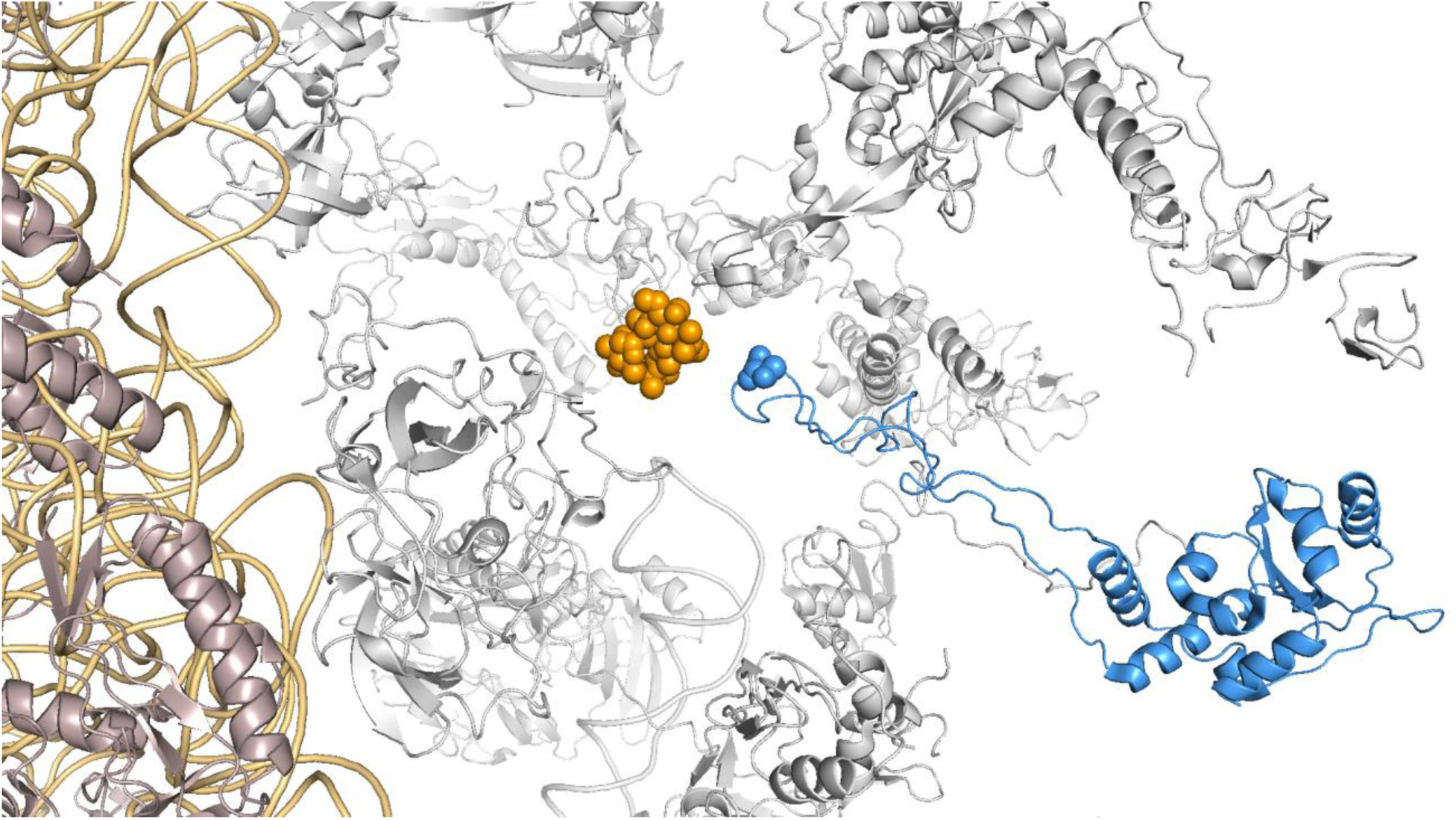
RplD G70 and the azithromycin binding pocket in the 50S ribosome from *Thermus thermophilus* (PDB ID: 4v7y). *N. gonorrhoeae* RplD is relatively similar to its *T. thermophilus* homolog (28.4% identical, 49.1% similarity using a BLOSUM62 matrix over 218 amino acids with 20 insertions/deletions). PyMOL (The PyMOL Molecular Graphics System, Version 2.0 Schrödinger, LLC) was used to depict azithromycin in orange and RplD in blue (with the G70 amino acid highlighted as blue spheres) and to hide the 23S rRNA for clarity.

**Supplementary Figure 5.**
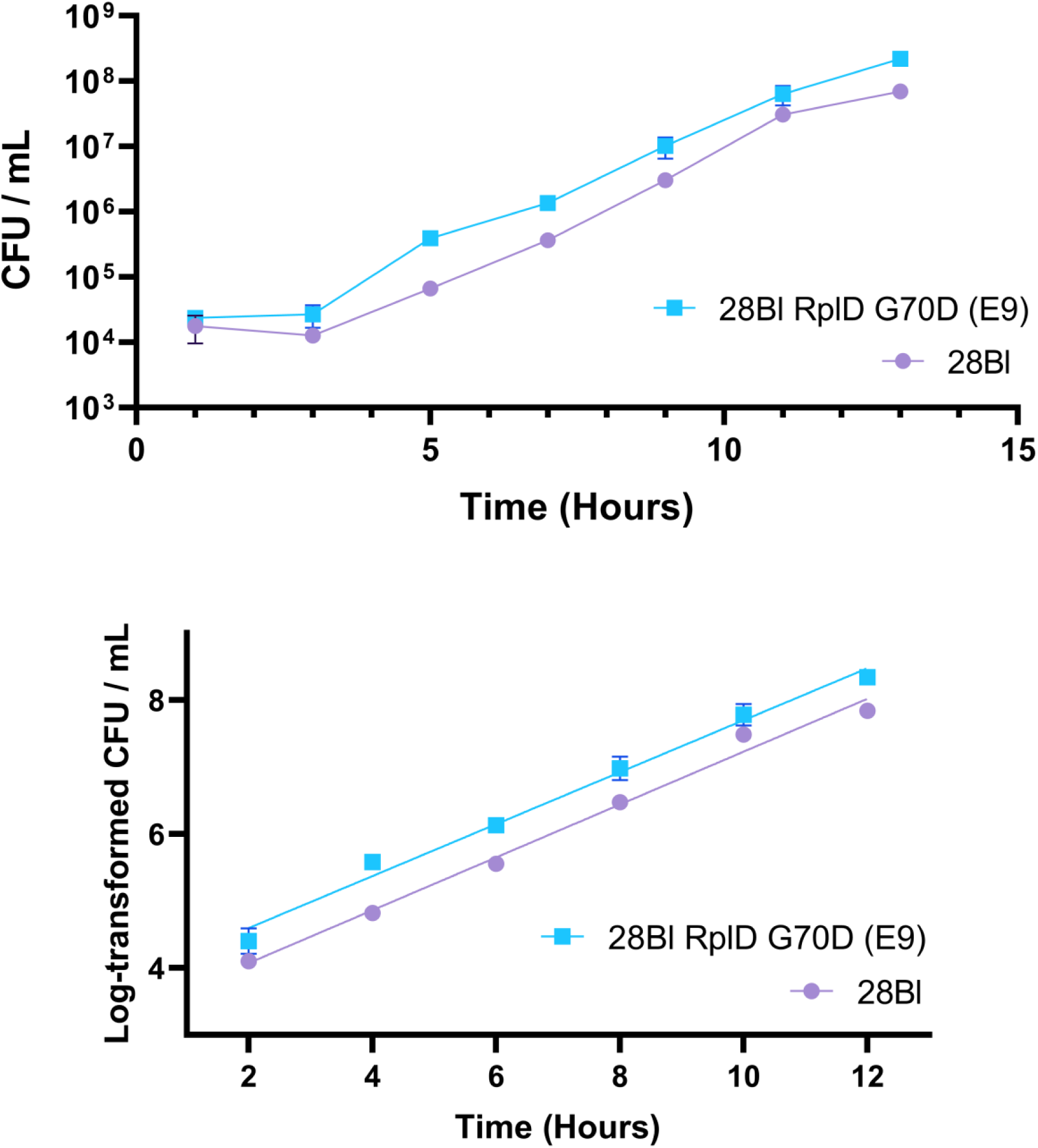
Growth curve experiments for RplD G70D isogenic strains. Error bars are SD for three technical replicates. Top – Calculated CFU / mLs from the full growth experiment for the two strains graphed on a logarithmic axis. Bottom – estimation of exponential phase best fit lines using GraphPad Prism following removal of lag phase data points and log-transformation of CFU / mLs; see Supplementary Table 5 for estimated parameters.

**Supplementary Table 4.** Prevalence of RplD macrolide binding site mutations across included datasets. Datasets with prevalence over 10% are highlighted.

| Dataset | Region and Timespan | Prevalence of RplD macrolide binding site mutations |
| --- | --- | --- |
| Demczuk et al., 2015 [59] | Canada 1989-2013 | 0.0439 |
| Demczuk et al., 2016 [37] | Canada 1982-2011 | 0.0804 |
| Eyre et al., 2017 [60] | Brighton, UK 2004-2011 | 0.0346 |
| Ezewudo et al., 2015 [61] | Global 1982-2011 | 0 |
| Fifer et al., 2018 [62] | UK 2004-2017 | 0.0200 |
| Grad et al., 2016 [3] and 2014 [63] | US 2000-2013 | 0.0582 |
| Harris et al., 2018 [64] | Europe 2013 | 0.0258 |
| Kwong et al., 2017 [65] | Melbourne, Australia 2005-2014 | 0.0532 |
| Lee et al., 2018 [66] | New Zealand 2014-2015 | 0.0050 |
| **Mortimer et al., 2020** | **NYC 2011-2015** | **0.1013** |
| Ryan et al., 2018 [67] | Ireland 2012-2016 | 0.0513 |
| Sánchez-Busó et al., 2019 [68] | Global 1979-2012 | 0.0212 |
| **Yahara et al., 2018 [27]** | **Kyoto and Osaka, Japan 1996-2015** | **0.1346** |

**Supplementary Table 5.**
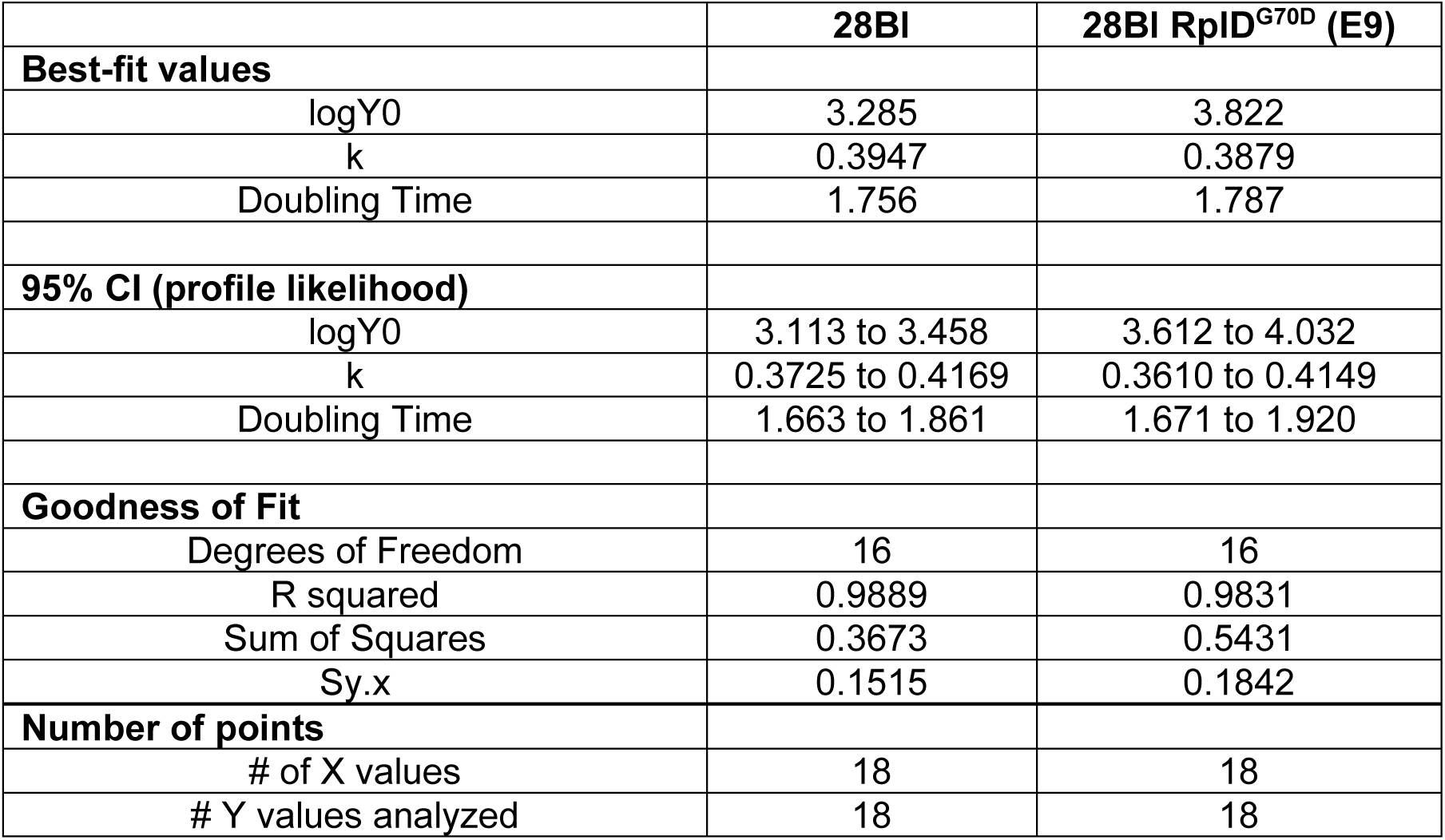
Estimated growth curve parameters. Estimation of exponential phase growth parameters using GraphPad Prism following removal of lag phase data points and log-transformation of CFU / mLs; see Supplementary Figure 6 (bottom) for estimated best fit lines.

## References

1. Unemo M, Shafer WM. Antimicrobial resistance in Neisseria gonorrhoeae in the 21st century: past, evolution, and future. Clin Microbiol Rev. 2014;27(3):587–613. Epub 2014/07/02. doi: 10.1128/CMR.00010-14. PubMed PMID: 24982323; PubMed Central PMCID: PMCPMC4135894.

2. Wi T, Lahra MM, Ndowa F, Bala M, Dillon JR, Ramon-Pardo P, et al. Antimicrobial resistance in Neisseria gonorrhoeae: Global surveillance and a call for international collaborative action. PLoS Med. 2017;14(7):e1002344. Epub 2017/07/08. doi: 10.1371/journal.pmed.1002344. PubMed PMID: 28686231; PubMed Central PMCID: PMCPMC5501266.

3. Grad YH, Harris SR, Kirkcaldy RD, Green AG, Marks DS, Bentley SD, et al. Genomic Epidemiology of Gonococcal Resistance to Extended-Spectrum Cephalosporins, Macrolides, and Fluoroquinolones in the United States, 2000-2013. J Infect Dis. 2016;214(10):1579–87. Epub 2016/10/30. doi: 10.1093/infdis/jiw420. PubMed PMID: 27638945; PubMed Central PMCID: PMCPMC5091375.

4. Fifer H, Saunders J, Soni S, Sadiq ST, Fitzgerald M. British Association for Sexual Health and HIV national guideline for the management of infection with Neisseria gonorrhoeae: BASHH; 2019. Available from: https://www.bashhguidelines.org/media/1208/gc-2019.pdf.

5. Allan-Blitz L-T, Hemarajata P, Humphries RM, Wynn A, Segura ER, Klausner JD. A Cost Analysis of Gyrase A Testing and Targeted Ciprofloxacin Therapy Versus Recommended 2-Drug Therapy for Neisseria gonorrhoeae Infection. Sex Transm Dis. 2018;45(2):87–91. doi: 10.1097/OLQ.0000000000000698.

6. Tuite AR, Gift TL, Chesson HW, Hsu K, Salomon JA, Grad YH. Impact of Rapid Susceptibility Testing and Antibiotic Selection Strategy on the Emergence and Spread of Antibiotic Resistance in Gonorrhea. J Infect Dis. 2017;216(9):1141–9. doi: 10.1093/infdis/jix450.

7. Ma KC, Mortimer TD, Hicks AL, Wheeler NE, Sánchez-Busó L, Golparian D, et al. Increased antibiotic susceptibility in Neisseria gonorrhoeae through adaptation to the cervical environment. bioRxiv. 2020. doi: 10.1101/2020.01.07.896696.

8. Hicks AL, Wheeler N, Sanchez-Buso L, Rakeman JL, Harris SR, Grad YH. Evaluation of parameters affecting performance and reliability of machine learning-based antibiotic susceptibility testing from whole genome sequencing data. PLoS Comput Biol. 2019;15(9):e1007349. Epub 2019/09/04. doi: 10.1371/journal.pcbi.1007349. PubMed PMID: 31479500; PubMed Central PMCID: PMCPMC6743791 that no competing interests exist.

9. Gaynor M, Mankin AS. Macrolide antibiotics: binding site, mechanism of action, resistance. Curr Top Med Chem. 2003;3(9):949–61. Epub 2003/04/08. doi: 10.2174/1568026033452159. PubMed PMID: 12678831.

10. Ng LK, Martin I, Liu G, Bryden L. Mutation in 23S rRNA associated with macrolide resistance in Neisseria gonorrhoeae. Antimicrob Agents Chemother. 2002;46(9):3020–5. Epub 2002/08/17. doi: 10.1128/aac.46.9.3020-3025.2002. PubMed PMID: 12183262; PubMed Central PMCID: PMCPMC127397.

11. Zhang J, van der Veen S. Neisseria gonorrhoeae 23S rRNA A2059G mutation is the only determinant necessary for high-level azithromycin resistance and improves in vivo biological fitness. J Antimicrob Chemother. 2019;74(2):407–15. doi: 10.1093/jac/dky438.

12. Rouquette C, Harmon JB, Shafer WM. Induction of the mtrCDE-encoded efflux pump system of Neisseria gonorrhoeae requires MtrA, an AraC-like protein. Mol Microbiol. 1999;33(3):651–8. Epub 1999/07/27. doi: 10.1046/j.1365-2958.1999.01517.x. PubMed PMID: 10417654.

13. Shafer WM, Folster JP, Warner DEM, Johnson PJT, Balthazar JT, Kamal N, et al. Expression of the MtrC-MtrD-MtrE Efflux Pump in Neisseria gonorrhoeae and Bacterial Survival in the Presence of Antimicrobials. National Institute of Allergy and Infectious Diseases, NIH; 2008: Humana Press; 2008. p. 55–63.

14. Veal WL, Nicholas RA, Shafer WM. Overexpression of the MtrC-MtrD-MtrE efflux pump due to an mtrR mutation is required for chromosomally mediated penicillin resistance in Neisseria gonorrhoeae. J Bacteriol. 2002;184(20):5619–24. Epub 2002/09/25. doi: 10.1128/jb.184.20.5619-5624.2002. PubMed PMID: 12270819; PubMed Central PMCID: PMCPMC139619.

15. Warner DM, Folster JP, Shafer WM, Jerse AE. Regulation of the MtrC-MtrD-MtrE efflux-pump system modulates the in vivo fitness of Neisseria gonorrhoeae. J Infect Dis. 2007;196(12):1804–12. Epub 2008/01/15. doi: 10.1086/522964. PubMed PMID: 18190261.

16. Warner DM, Shafer WM, Jerse AE. Clinically relevant mutations that cause derepression of the Neisseria gonorrhoeae MtrC-MtrD-MtrE Efflux pump system confer different levels of antimicrobial resistance and in vivo fitness. Mol Microbiol. 2008;70(2):462–78. Epub 2008/09/03. doi: 10.1111/j.1365-2958.2008.06424.x. PubMed PMID: 18761689; PubMed Central PMCID: PMCPMC2602950.

17. Zalucki YM, Dhulipala V, Shafer WM. Dueling regulatory properties of a transcriptional activator (MtrA) and repressor (MtrR) that control efflux pump gene expression in Neisseria gonorrhoeae. MBio. 2012;3(6):e00446–12. doi: 10.1128/mBio.00446-12. PubMed Central PMCID: PMCPMC3517864.

18. Zarantonelli L, Borthagaray G, Lee E-H, Shafer WM. Decreased Azithromycin Susceptibility ofNeisseria gonorrhoeae Due to mtrRMutations. Antimicrob Agents Chemother. 1999;43(10):2468–72.

19. Rouquette-Loughlin CE, Reimche JL, Balthazar JT, Dhulipala V, Gernert KM, Kersh EN, et al. Mechanistic Basis for Decreased Antimicrobial Susceptibility in a Clinical Isolate of Neisseria gonorrhoeae Possessing a Mosaic-Like mtr Efflux Pump Locus. MBio. 2018;9(6). Epub 2018/11/30. doi: 10.1128/mBio.02281-18. PubMed PMID: 30482834; PubMed Central PMCID: PMCPMC6282211.

20. Wadsworth CB, Arnold BJ, Sater MRA, Grad YH. Azithromycin Resistance through Interspecific Acquisition of an Epistasis-Dependent Efflux Pump Component and Transcriptional Regulator in Neisseria gonorrhoeae. MBio. 2018;9(4). Epub 2018/08/09. doi: 10.1128/mBio.01419-18. PubMed PMID: 30087172; PubMed Central PMCID: PMCPMC6083905.

21. Cousin SL, Jr., Whittington WLH, Roberts MC. Acquired macrolide resistance genes and the 1 bp deletion in the mtrR promoter in Neisseria gonorrhoeae. J Antimicrob Chemother. 2003;51(1):131–3.

22. Diner EJ, Hayes CS. Recombineering reveals a diverse collection of ribosomal proteins L4 and L22 that confer resistance to macrolide antibiotics. J Mol Biol. 2009;386(2):300–15. doi: 10.1016/j.jmb.2008.12.064. PubMed PMID: 19150357; PubMed Central PMCID: PMCPMC2644216.

23. Wheeler NE, Reuter S, Chewapreecha C, Lees JA, Blane B, Horner C, et al. Contrasting approaches to genome-wide association studies impact the detection of resistance mechanisms in Staphylococcus aureus. bioRxiv. 2019. doi: 10.1101/758144.

24. Belkacem A, Jacquier H, Goubard A, Mougari F, La Ruche G, Patey O, et al. Molecular epidemiology and mechanisms of resistance of azithromycin-resistant Neisseria gonorrhoeae isolated in France during 2013-14. J Antimicrob Chemother. 2016;71(9):2471–8. doi: 10.1093/jac/dkw182. PubMed PMID: 27301565.

25. Jacobsson S, Golparian D, Cole M, Spiteri G, Martin I, Bergheim T, et al. WGS analysis and molecular resistance mechanisms of azithromycin-resistant (MIC >2 mg/L) Neisseria gonorrhoeae isolates in Europe from 2009 to 2014. J Antimicrob Chemother. 2016;71(11):3109–16. doi: 10.1093/jac/dkw279. PubMed PMID: 27432597.

26. Demczuk W, Martin I, Sawatzky P, Allen V, Lefebvre B, Hoang L, et al. Equations to predict antimicrobial minimum inhibitory concentrations in Neisseria gonorrhoeae using molecular antimicrobial resistance determinants. Antimicrob Agents Chemother. 2019. doi: 10.1128/AAC.02005-19.

27. Yahara K, Nakayama SI, Shimuta K, Lee KI, Morita M, Kawahata T, et al. Genomic surveillance of Neisseria gonorrhoeae to investigate the distribution and evolution of antimicrobial-resistance determinants and lineages. Microb Genom. 2018;4(8). Epub 2018/08/01. doi: 10.1099/mgen.0.000205. PubMed PMID: 30063202; PubMed Central PMCID: PMCPMC6159555.

28. Hicks AL, Kissler SM, Mortimer TD, Ma KC, Taiaroa G, Ashcroft M, et al. Targeted surveillance strategies for efficient detection of novel antibiotic resistance variants. bioRxiv. 2020.

29. Zheng Z, Liu L, Shen X, Yu J, Chen L, Zhan L, et al. Antimicrobial Resistance And Molecular Characteristics Among Neisseria gonorrhoeae Clinical Isolates In A Chinese Tertiary Hospital. Infect Drug Resist. 2019;12:3301–9. doi: 10.2147/IDR.S221109. PubMed PMID: 31695449; PubMed Central PMCID: PMCPMC6815782.

30. Zeitouni S, Collin O, Andraud M, Ermel G, Kempf I. Fitness of macrolide resistant Campylobacter coli and Campylobacter jejuni. Microb Drug Resist. 2012;18(2):101–8. doi: 10.1089/mdr.2011.0188.

31. Saber M, Shapiro B. Benchmarking bacterial genome-wide association study methods using simulated genomes and phenotypes. Microbial Genomics. 2020. doi: doi:10.1099/mgen.0.000337.

32. Ma L, Han S, Yang J, Da Y. Multi-locus test conditional on confirmed effects leads to increased power in genome-wide association studies. PLoS One. 2010;5(11):e15006. Epub 2010/11/26. doi: 10.1371/journal.pone.0015006. PubMed PMID: 21103364; PubMed Central PMCID: PMCPMC2982824.

33. Segura V, Vilhjalmsson BJ, Platt A, Korte A, Seren U, Long Q, et al. An efficient multilocus mixed-model approach for genome-wide association studies in structured populations. Nat Genet. 2012;44(7):825–30. Epub 2012/06/19. doi: 10.1038/ng.2314. PubMed PMID: 22706313; PubMed Central PMCID: PMCPMC3386481.

34. Lees JATM, T; Galardini, Marco; Wheeler, Nicole E; Corander, Jukka. Improved inference and prediction of bacterial genotype-phenotype associations using pangenome-spanning regressions. bioRxiv. 2019. doi: 10.1101/852426.

35. Thomas JC, Seby S, Abrams AJ, Cartee J, Lucking S, Vidyaprakash E, et al. Evidence of Recent Genomic Evolution in Gonococcal Strains with Decreased Susceptibility to Cephalosporins or Azithromycin in the United States, 2014-2016. J Infect Dis. 2019. doi: 10.1093/infdis/jiz079.

36. Golparian D, Harris SR, Sanchez-Buso L, Hoffmann S, Shafer WM, Bentley SD, et al. Genomic evolution of Neisseria gonorrhoeae since the preantibiotic era (1928-2013): antimicrobial use/misuse selects for resistance and drives evolution. BMC Genomics. 2020;21(1):116. doi: 10.1186/s12864-020-6511-6. PubMed PMID: 32013864; PubMed Central PMCID: PMCPMC6998845.

37. Demczuk W, Martin I, Peterson S, Bharat A, Van Domselaar G, Graham M, et al. Genomic Epidemiology and Molecular Resistance Mechanisms of Azithromycin-Resistant Neisseria gonorrhoeae in Canada from 1997 to 2014. J Clin Microbiol. 2016;54(5):1304–13. Epub 2016/03/05. doi: 10.1128/JCM.03195-15. PubMed PMID: 26935729; PubMed Central PMCID: PMCPMC4844716.

38. Endimiani A, Guilarte YN, Tinguely R, Hirzberger L, Selvini S, Lupo A, et al. Characterization of Neisseria gonorrhoeae isolates detected in Switzerland (1998-2012): emergence of multidrug-resistant clones less susceptible to cephalosporins. BMC Infect Dis. 2014;14:106. doi: 10.1186/1471-2334-14-106. PubMed PMID: 24568221; PubMed Central PMCID: PMCPMC3941690.

39. Wan C, Li Y, Le W-J, Liu Y-R, Li S, Wang B, et al. Increasing resistance to azithromycin ofNeisseria gonorrhoeaein eastern Chinese cities: mechanisms and genetic diversity of resistant Nanjing isolates. Antimicrob Agents Chemother. 2018. doi: 10.1128/AAC.02499-17.

40. Harrison OB, Clemence M, Dillard JP, Tang CM, Trees D, Grad YH, et al. Genomic analyses of Neisseria gonorrhoeae reveal an association of the gonococcal genetic island with antimicrobial resistance. J Infect. 2016;73(6):578–87. doi: 10.1016/j.jinf.2016.08.010. PubMed PMID: 27575582; PubMed Central PMCID: PMCPMC5127880.

41. Colangeli R, Jedrey H, Kim S, Connell R, Ma S, Chippada Venkata UD, et al. Bacterial Factors That Predict Relapse after Tuberculosis Therapy. N Engl J Med. 2018;379(9):823–33. Epub 2018/08/30. doi: 10.1056/NEJMoa1715849. PubMed PMID: 30157391; PubMed Central PMCID: PMCPMC6317071.

42. Yasuda M, Ito S, Hatazaki K, Deguchi T. Remarkable increase of Neisseria gonorrhoeae with decreased susceptibility of azithromycin and increase in the failure of azithromycin therapy in male gonococcal urethritis in Sendai in 2015. J Infect Chemother. 2016;22(12):841–3. Epub 2016/09/01. doi: 10.1016/j.jiac.2016.07.012. PubMed PMID: 27578029.

43. Tapsall JW, Shultz TR, Limnios EA, Donovan B, Lum G, Mulhall BP. Failure of azithromycin therapy in gonorrhea and discorrelation with laboratory test parameters. Sex Transm Dis. 1998;25(10):505–8. doi: 10.1097/00007435-199811000-00002.

44. Fifer H, Cole M, Hughes G, Padfield S, Smolarchuk C, Woodford N, et al. Sustained transmission of high-level azithromycin-resistant Neisseria gonorrhoeae in England: an observational study. Lancet Infect Dis. 2018. doi: 10.1016/S1473-3099(18)30122-1.

45. Croucher NJ, Page AJ, Connor TR, Delaney AJ, Keane JA, Bentley SD, et al. Rapid phylogenetic analysis of large samples of recombinant bacterial whole genome sequences using Gubbins. Nucleic Acids Res. 2015;43(3):e15. Epub 2014/11/22. doi: 10.1093/nar/gku1196. PubMed PMID: 25414349; PubMed Central PMCID: PMCPMC4330336.

46. Walker BJ, Abeel T, Shea T, Priest M, Abouelliel A, Sakthikumar S, et al. Pilon: an integrated tool for comprehensive microbial variant detection and genome assembly improvement. PLoS One. 2014;9(11):e112963. Epub 2014/11/20. doi: 10.1371/journal.pone.0112963. PubMed PMID: 25409509; PubMed Central PMCID: PMCPMC4237348.

47. Li H. Aligning sequence reads, clone sequences and assembly contigs with BWA-MEM. arXiv preprint. 2013;1303.3997.

48. Lees JA, Galardini M, Bentley SD, Weiser JN, Corander J. pyseer: a comprehensive tool for microbial pangenome-wide association studies. Bioinformatics. 2018;34(24):4310–2. Epub 2018/12/12. doi: 10.1093/bioinformatics/bty539. PubMed PMID: 30535304; PubMed Central PMCID: PMCPMC6289128.

49. Bankevich A, Nurk S, Antipov D, Gurevich AA, Dvorkin M, Kulikov AS, et al. SPAdes: a new genome assembly algorithm and its applications to single-cell sequencing. J Comput Biol. 2012;19(5):455–77. Epub 2012/04/18. doi: 10.1089/cmb.2012.0021. PubMed PMID: 22506599; PubMed Central PMCID: PMCPMC3342519.

50. Kersh EN, Allen V, Ransom E, Schmerer M, St Cyr S, Workowski K, et al. Rationale for a Neisseria gonorrhoeae Susceptible Only Interpretive Breakpoint for Azithromycin. Clin Infect Dis. 2019. Epub 2019/04/10. doi: 10.1093/cid/ciz292. PubMed PMID: 30963175; PubMed Central PMCID: PMCPMC6785360.

51. Letunic I, Bork P. Interactive Tree Of Life (iTOL) v4: recent updates and new developments. Nucleic Acids Res. 2019;47(W1):W256-W9. Epub 2019/04/02. doi: 10.1093/nar/gkz239. PubMed PMID: 30931475; PubMed Central PMCID: PMCPMC6602468.

52. Arnold B, Sohail M, Wadsworth C, Corander J, Hanage WP, Sunyaev S, et al. Finescale haplotype structure reveals strong signatures of positive selection in a recombining bacterial pathogen. Mol Biol Evol. 2019. doi: 10.1093/molbev/msz225.

53. Hill WG, Robertson A. Linkage disequilibrium in finite populations. Theor Appl Genet. 1968;38(6):226–31. doi: 10.1007/BF01245622. PubMed PMID: 24442307.

54. Altschul SF, Gish W, Miller W, Myers EW, Lipman DJ. Basic local alignment search tool. J Mol Biol. 1990;215(3):403–10. Epub 1990/10/05. doi: 10.1016/S0022-2836(05)80360-2. PubMed PMID: 2231712.

55. Katoh K, Standley DM. MAFFT multiple sequence alignment software version 7: improvements in performance and usability. Mol Biol Evol. 2013;30(4):772–80. doi: 10.1093/molbev/mst010. PubMed Central PMCID: PMCPMC3603318.

56. Dillard JP. Genetic Manipulation of Neisseria gonorrhoeae. Curr Protoc Microbiol. 2011;Chapter 4:Unit4A 2. Epub 2011/11/03. doi: 10.1002/9780471729259.mc04a02s23. PubMed PMID: 22045584; PubMed Central PMCID: PMCPMC4549065.

57. Ambur OH, Frye SA, Tønjum T. New functional identity for the DNA uptake sequence in transformation and its presence in transcriptional terminators. J Bacteriol. 2007;189(5):2077–85. doi: 10.1128/JB.01408-06. PubMed Central PMCID: PMCPMC1855724.

58. Wade JJ, Graver MA. A fully defined, clear and protein-free liquid medium permitting dense growth of Neisseria gonorrhoeae from very low inocula. FEMS Microbiol Lett. 2007;273(1):35–7. doi: 10.1111/j.1574-6968.2007.00776.x.

59. Demczuk W, Lynch T, Martin I, Van Domselaar G, Graham M, Bharat A, et al. Wholegenome phylogenomic heterogeneity of Neisseria gonorrhoeae isolates with decreased cephalosporin susceptibility collected in Canada between 1989 and 2013. J Clin Microbiol. 2015;53(1):191–200. Epub 2014/11/08. doi: 10.1128/JCM.02589-14. PubMed PMID: 25378573; PubMed Central PMCID: PMCPMC4290921.

60. Eyre DW, De Silva D, Cole K, Peters J, Cole MJ, Grad YH, et al. WGS to predict antibiotic MICs for Neisseria gonorrhoeae. J Antimicrob Chemother. 2017;72(7):1937–47. Epub 2017/03/24. doi: 10.1093/jac/dkx067. PubMed PMID: 28333355; PubMed Central PMCID: PMCPMC5890716.

61. Ezewudo MN, Joseph SJ, Castillo-Ramirez S, Dean D, Del Rio C, Didelot X, et al. Population structure of Neisseria gonorrhoeae based on whole genome data and its relationship with antibiotic resistance. PeerJ. 2015;3:e806. Epub 2015/03/18. doi: 10.7717/peerj.806. PubMed PMID: 25780762; PubMed Central PMCID: PMCPMC4358642.

62. Fifer H, Cole M, Hughes G, Padfield S, Smolarchuk C, Woodford N, et al. Sustained transmission of high-level azithromycin-resistant Neisseria gonorrhoeae in England: an observational study. Lancet Infect Dis. 2018;18(5):573–81. Epub 2018/03/11. doi: 10.1016/S1473-3099(18)30122-1. PubMed PMID: 29523496.

63. Grad YH, Kirkcaldy RD, Trees D, Dordel J, Harris SR, Goldstein E, et al. Genomic epidemiology of Neisseria gonorrhoeae with reduced susceptibility to cefixime in the USA: a retrospective observational study. Lancet Infect Dis. 2014;14(3):220–6. Epub 2014/01/28. doi: 10.1016/S1473-3099(13)70693-5. PubMed PMID: 24462211; PubMed Central PMCID: PMCPMC4030102.

64. Harris SR, Cole MJ, Spiteri G, Sanchez-Buso L, Golparian D, Jacobsson S, et al. Public health surveillance of multidrug-resistant clones of Neisseria gonorrhoeae in Europe: a genomic survey. Lancet Infect Dis. 2018;18(7):758–68. Epub 2018/05/20. doi: 10.1016/S1473-3099(18)30225-1. PubMed PMID: 29776807; PubMed Central PMCID: PMCPMC6010626.

65. Kwong JC, Chow EPF, Stevens K, Stinear TP, Seemann T, Fairley CK, et al. Whole-genome sequencing reveals transmission of gonococcal antibiotic resistance among men who have sex with men: an observational study. Sex Transm Infect. 2018;94(2):151–7. doi: 10.1136/sextrans-2017-053287. PubMed PMID: 29247013; PubMed Central PMCID: PMCPMC5870456.

66. Lee RS, Seemann T, Heffernan H, Kwong JC, Goncalves da Silva A, Carter GP, et al. Genomic epidemiology and antimicrobial resistance of Neisseria gonorrhoeae in New Zealand. J Antimicrob Chemother. 2018;73(2):353–64. doi: 10.1093/jac/dkx405. PubMed PMID: 29182725; PubMed Central PMCID: PMCPMC5890773.

67. Ryan L, Golparian D, Fennelly N, Rose L, Walsh P, Lawlor B, et al. Antimicrobial resistance and molecular epidemiology using whole-genome sequencing of Neisseria gonorrhoeae in Ireland, 2014-2016: focus on extended-spectrum cephalosporins and azithromycin. Eur J Clin Microbiol Infect Dis. 2018. doi: 10.1007/s10096-018-3296-5. PubMed PMID: 29882175.

68. Sanchez-Buso L, Golparian D, Corander J, Grad YH, Ohnishi M, Flemming R, et al. The impact of antimicrobials on gonococcal evolution. Nat Microbiol. 2019. Epub 2019/07/31. doi: 10.1038/s41564-019-0501-y. PubMed PMID: 31358980.

